# Sub-chronic peripheral CB1R inhibition enhances cognitive performance and induces hippocampal synaptic plasticity changes in naïve mice

**DOI:** 10.1101/2023.11.21.567995

**Authors:** Araceli Bergadà-Martínez, Lucía de los Reyes-Ramírez, Sara Martínez-Torres, Irene Martínez- Gallego, Rafael Maldonado, Antonio Rodríguez-Moreno, Andrés Ozaita

**Affiliations:** Laboratory of Neuropharmacology-NeuroPhar, Department of Medicine and Life Sciences, Universitat Pompeu Fabra, Barcelona, Spain; Laboratory of Cellular Neuroscience and Plasticity, Department of Physiology, Anatomy and Cell Biology, University Pablo de Olavide, Seville, Spain; IMIM Hospital del Mar Research Institute, Barcelona, Spain

**Author notes:** Corresponding author: Andrés Ozaita, Laboratori de Neurofarmacologia, Facultat de Ciències de la Salut i de la Vida, Universitat Pompeu Fabra, Parc de Recerca Biomèdica de Barcelona, C/Doctor Aiguader 88, 08003 Barcelona, Spain.

**Keywords:** peripheral-to-central, cannabinoid type-1 receptor, cognitive enhancement, hippocampal synaptic plasticity

## Abstract

**Background and Purpose:** The peripheral contribution to brain function and cognitive performance is far from understood. Cannabinoid type-1 receptor (CB1R) is classically pictured in the central nervous system to have such a role. We previously demonstrated a novel mechanism where the acute peripheral CB1R inhibition in mice prolongs memory persistence. Here, we take advance of the repeated exposure to the peripherally-restricted CB1R antagonist to further reveal cognitive improvements and the hippocampal mechanisms involved.

**Experimental Approach:** We evaluated in young adult male and female mice the behavioural consequences of a sub-chronic treatment with AM6545. Moreover, an unbiased transcriptomic analysis, as well as electrophysiological and biochemical studies, were performed in the hippocampus of treated mice to elucidate the central cellular and molecular consequences of such peripheral approach.

**Key Results:** Sub-chronic inhibition of peripheral CB1R with AM6545 resulted in enhanced memory in low and high arousal conditions. Moreover, executive function was facilitated after repeated AM6545 administration, further strengthening the cognitive improving properties of peripheral CB1R inhibition. Transcriptional analysis of hippocampal synaptoneurosomes from treated male and female mice revealed a sex-dependent modulation of synaptic transcripts by AM6545. Notably, AM6545 occluded long-term potentiation in CA3-CA1 synapses while enhancing input-output relation. This was accompanied by an increase in the hippocampal expression of *Bdnf* and *Ngf*.

**Conclusion and Implications:** Our results show that peripheral CB1R inhibition contributes to the modulation of memory persistence, executive function, and hippocampal synaptic plasticity in mice, further indicating that peripheral CB1R could act as a target for a novel class of nootropic compounds.

**What is already known?:**

- Acute peripheral CB1R inhibition enhances object-recognition memory persistence in mice.
- Such enhancement occurs through a noradrenergic mechanism involving the vagus nerve.

**What does this study add?:**

- Peripheral CB1R inhibition modifies synaptoneurosomal transcriptome in the hippocampus of mice.
- No tolerance and no side effects were observed after peripheral CB1R inhibition.

**What is the clinical significance?:**

- Peripheral CB1R inhibition may function as a novel strategy for cognitive improvement.

## Introduction

Most everyday events create low arousal non-emotional memory traces which may fade away or persist through unpredictable periods that may go on for as long as a lifetime (Morris, 2006). The persistence of memory is facilitated in emotional memories, where encoding is intermingled with the natural stress-coping response, leading to a significant enhancement in memory recall for the sensory information encoded at that time (Roozendaal et al., 2009).

The endocannabinoid system (ECS) is an endogenous neuromodulatory system highly expressed in the central nervous system (CNS) and peripheral tissues, and plays a relevant role in modulating learning and memory processes. Cannabinoid type-1 receptor (CB1R) is the most widely distributed G-protein coupled receptor subtype in the CNS. Within the brain, CB1R is highly expressed in regions such as the hippocampus or the cortex, as well as other memory-related regions (Pacher et al., 2006; Zanettini et al., 2011). Besides its high expression in the brain, CB1R is also present in many peripheral tissues such as the liver, adipose tissue, endocrine system, or skeletal muscle. Regarding its subcellular distribution, CB1R is detected predominantly at presynaptic sites in neurons where it suppresses neurotransmitter/neuromodulator release depending on local synaptic activity (Ohno-Shosaku and Kano, 2014). For example, activation of the CB1R in adrenergic and noradrenergic cells leads to a decrease in the release of adrenaline and noradrenaline highlighting the relevance of CB1R in neurotransmitter/neuromodulator release function (Ishac et al., 1996; Niederhoffer et al., 2001). Notably, the administration of CB1R agonists in rodents results in deficits in several memory types such as recognition memory (Puighermanal et al., 2009), spatial memory (Da Silva and Takahashi, 2002) or fear memory (Pamplona and Takahashi, 2006; Kruk-Slomka and Biala, 2016). On the contrary, CB1R antagonists prevented some of these memory impairments (Pamplona and Takahashi, 2006; Barna et al., 2007). Interestingly, our group previously demonstrated that the acute inhibition of peripheral CB1R triggers an adrenergic mechanism that enhances memory persistence involving the adrenal glands and the vagus nerve. We found this acute cognitive enhancement is accompanied by changes in functional brain connectivity as well as an increase in hippocampal extracellular noradrenaline (Martínez-Torres et al., 2023).

In the present study, we used AM6545 (Tam et al., 2010), the peripheral antagonist of CB1R, to reveal modulation in different cognitive domains, including non-emotional (low arousal) and emotional (high arousal) memory and executive function. We also explored the synaptic mechanisms involved in cognitive facilitation. For this purpose, we combined behavioural tests with transcriptomic, electrophysiological and biochemical studies. We found that sub-chronic peripheral CB1R inhibition results in enhanced cognitive performance in both male and female mice. Moreover, we report a sex-dependent modulation in synaptic transcripts, and synaptic plasticity modifications in the hippocampus, including modulation of long-term potentiation and enhanced expression of neurotrophic factors. These central effects of repeated peripheral CB1R inhibition unmask a target worth exploring in the context of cognitive improvement.

## Materials and methods

### Ethics

All animal procedures were conducted following “Animals in Research: Reporting Experiments” (ARRIVE) guidelines and standard ethical guidelines (Kilkenny et al., 2010) (European Directive 2010/63/EU). Animal procedures were approved by the local ethical committee (Comitè Ètic d’Experimentació Animal-Parc de Recerca Biomèdica de Barcelona, CEEA-PRBB).

### Animals

Young adult (8-12 weeks old) male and female Swiss albino (CD-1) and C57BL/6J mice (Charles River) were used for pharmacological approaches on behavioural, transcriptomic, electrophysiological and biochemical experiments. Mice were housed in Plexiglas cages (2-4 mice per cage in the case of males and 2-5 in the case of females) and maintained in temperature (21 ± 1 °C) and humidity (55 ± 10%) controlled environment. Food and water were available *ad libitum*. Animals tested in the touchscreen apparatus were food-restricted to 85-95% of their free-feeding weight beginning 3 days before pre-training. These mice were maintained on food restriction for the entire experiment although daily food allowance was increased over time to prevent stunting. All the experiments were performed during the light phase of a 12 h cycle (light on at 8 am; light off at 8 pm). Mice were habituated to the experimental room and handled for 1 week before starting the experiments. All behavioural experiments were conducted by an observer blind to experimental conditions.

### Drugs and treatments

AM6545 (Tocris-Bio-Techne) was dissolved in 0.26% DMSO, 4.74% ethanol, 5% cremophor-EL and 90% saline. Sotalol (Sigma-Aldrich) was dissolved in saline (Martínez-Torres et al., 2023). All drugs were injected intraperitoneally (i.p.) in a volume of 10 mL/kg of body weight. AM6545 was administered at the dose of 1 mg/kg and sotalol at the dose of 10 mg/kg (Martínez-Torres et al., 2023). Sub-chronic administrations consisted of 7 days of treatment. For the pairwise discrimination acquisition, AM6545 was administered 30 min after the end of the pairwise discrimination session every day until the experiment finished.

### Novel object-recognition test

Novel object-recognition test (NORT) was performed as previously described (Martínez-Torres et al., 2023). Briefly, on day 1 mice were habituated to a V-shaped maze (V-maze) for 9 min. On day 2 (training phase), two identical objects (familiar objects) were located at the end of each corridor and mice were left to explore them for 9 min. To assess memory persistence, the test phase, also of 9 min, was performed 48 h later, and one of the familiar objects was replaced by a new object (novel object). The time spent exploring each of the objects was computed to calculate a discrimination index (DI = (Time Novel Object - Time Familiar Object)/(Time Novel Object + Time Familiar Object)). Object exploration time was considered when the orientation of the nose was towards the object. A higher discrimination index is considered to reflect greater memory retention for the familiar object. Importantly, in this work a variation of this protocol was applied when required, which consisted of a reduction in the training phase to 3 min and performing the test phase 24 h later. These two versions of the task allow assessing memory under challenging conditions. For this task, drug treatment was administered after the training phase in sub-chronic treatments. For the measurement of the effects after treatment withdrawal the treatment finished 24 h prior to the training phase.

### Novel place-recognition test

On days 1 and 2 mice were introduced into an empty arena (25×25×15 cm) with spatial cues and dimly illuminated (2-3 lux) and were allowed to freely explore it during 9 min (habituation phase). On day 3, two identical objects were positioned in opposite corners (familiar location) (6 cm away from the walls) and mice were allowed to explore them during 9 min (training phase). 48 h later, mice were introduced in the same arena with the same objects, but one of them was moved to a new location (novel location) (another corner, 6 cm away from the walls). Mice were allowed to explore the objects for 9 min (test phase). The time spent exploring each of the objects was computed to calculate a discrimination index (DI = (Time Novel Location - Time Familiar Location)/(Time Novel Location + Time Familiar Location)). Object exploration time was considered when the orientation of the nose was towards the object. A higher discrimination index is considered to reflect greater memory retention for the familiar location.

### Emotional memory persistence assessment in the context fear conditioning

Context fear conditioning (FC) was used to measure the effect of AM6545 on hippocampal-dependent emotional memory in mice. On day 1, mice were placed in a conditioning chamber with an electrifiable floor. After 3 min of free exploration, mice received one footshock (unconditioned stimulus, US: 2 s, 0.35 mA intensity) (training phase). On day 2 (first extinction session), 24 h after the training phase, mice were placed for 5 min in absence of the footshock in the same conditioning chamber and the freezing behaviour was manually recorded. Extinction sessions (Ext1-Ext5) were performed at 24, 48, 72, 96 and 120 h after the footshock, and freezing behaviour was manually counted. Treatment with AM6545 was administered immediately after each extinction session.

### Elevated plus maze test

The elevated plus maze test was performed as previously described (Navarro-Romero et al., 2022). The test consisted of a black Plexiglas apparatus with four arms (29 cm long × 5 cm wide) set in a cross from a neutral central square (5 cm × 5 cm) elevated 40 cm above the floor. Five-minute test sessions were performed after 7 days of treatment, 24 h after the last AM6545 or vehicle administration. The percentage of time and entries in the open arms were determined.

### Locomotor activity

Locomotor activity was assessed for 30 min after 7 days of AM6545 treatment 24 h after the last drug administration. Individual locomotor activity boxes (9 × 20 × 11 cm) (Imetronic) were used in a low luminosity environment as previously described (Navarro-Romero et al., 2022). The number of horizontal movements was detected by a line of photocells located 2 cm above the floor.

### Touchscreen apparatus

Automated touchscreen operant systems (Campden Instruments) were used to test learning (Horner et al., 2013). The system was controlled by ABET II software (Campden Instruments). Male mice began the experiment at 8-12 weeks of age and were tested in the same chamber daily five days a week until the end of the experiment.

### Pairwise discrimination task

Mice were habituated to the chamber prior to pre-training and testing. During habituation, the mouse was left in the chamber for 20 min. A tone was played and the reward magazine illuminated indicating the delivery of 10% condensed milk (CM) (150 μl). After a nosepoke there was a 10 s delay before a tone was played, the reward magazine illuminated again and CM was then delivered (7 μl). The procedure was repeated until the session ended. Following habituation, mice started the pre-training which consisted of four different stages. Each session lasted a maximum of 60 min. On stage 1, images/stimuli were displayed one at a time on one side of the touchscreen. The other side remained blank. Trials resulted in the delivery of the CM either upon touching the image or after an intertrial interval (ITI) of 20 s, whichever came first. On stage 2, the reinforcement was only delivered when mice touched the image. Mice remained in this stage until they performed 30 trials in a session. On stage 3, mice had to enter the reward magazine to initiate each trial. Mice remained in this stage until they performed 30 trials in a session. On stage 4, if mice touched the blank side of the touchscreen the house light illuminated for a time out period (5 s) and no reward was given. Mice remained in this stage until they performed 23 correct trials out of 30 in a session. Finally, in the pairwise discrimination acquisition, mice were taught to discriminate and choose a rewarded image (S+) over an unrewarded image (S-). The two images were randomly presented on the right or left side of the touchscreen. Touching the correct stimulus resulted in the delivery of the reward, whereas touching the incorrect stimulus resulted in the illumination of the house light. An incorrect trial led to a correction trial in which the two images were presented in the same location as in the previous trial. Correction trials were not included in the final correct trials count. Mice completed the task when they performed 24 correct trials out of 30 (80%) for two days in a row. Animals that did not achieve the criterion within 20 sessions were excluded from the experiment (Four animals were excluded, one vehicle-treated and three AM6545-treated).

### Tissue preparation for mRNA and biochemical analysis

Mice were treated with AM6545 (1 mg/kg, i.p.) or vehicle for 7 days. 24 h after the last administration, hippocampal tissues were dissected on ice, frozen on dry ice and stored at -80 °C until used.

### Tissue preparation for RNA extraction

*Synaptoneurosome preparation*: for RNA of a synaptoneurosome-enriched fraction, synaptoneurosome preparation was performed as previously described (Salgado-Mendialdúa et al., 2018). Hippocampal tissue was Dounce-homogenized by 10 strokes with a loose pestle and 10 strokes with a tight pestle in 30 volumes of ice-cold synaptoneurosome lysis buffer (in mM: 2.5 CaCl_2_, 124 NaCl, 3.2 KCl, 1.06 KH_2_PO, 26 NaHCO_3_, 1.3 MgCl_2_, 10 Glucose, 320 Sucrose, 20 HEPES/NaOH pH7.4) including phosphatase inhibitors (in mM: 5 NaPyro, 100 NaF, 1 NaOrth, 40 beta-glycerolphosphate) and protease inhibitors (1 μg/mL leupeptin, 10 μg/mL aprotinin, 1 μg/mL pepstatin and 1 mM phenylmethylsulfonyl fluoride). This crude homogenate was centrifuged for 1 min at 2,000 g, 4 °C, and the supernatant was recovered (S1). The pellet was resuspended in 1 mL of synaptoneurosome lysis buffer for further centrifugation (1 min at 2,000 g, 4 °C). This second supernatant (S2) was recovered and combined with S1. Total supernatant (S1 + S2) was passed through a 10 μm hydrophobic PTFE filter (LCWP02500, Merck Millipore) and centrifuged for 1 min at 4,000 g, 4 °C to attain the supernatant (S3). S3 was transferred to a new tube and centrifuged for 4 min at 14,000 g, 4 °C. This final supernatant (S4) was discarded, and the pellet was used as the synaptoneurosomal fraction. Subsequent RNA extraction was performed using a RNeasy Mini kit (74140, QIAGEN) according to the manufacturer’s instructions.

*Total homogenate*: for RNA of total hippocampal homogenates, hippocampal tissue was Dounce-homogenized using a RNeasy Mini kit (74140, QIAGEN) according to the manufacturer’s instructions.

### Hippocampal synaptoneurosomal RNA Sequencing

mRNA libraries from C57BL/6J male and female mice (VEH-male, *n* = 5; AM6545-male, *n* = 5; VEH-female, *n* = 3; AM6545-female, *n* = 4) were obtained by Macrogen Inc. using SMARTer Ultra Low Input kit (634940, Takara Bio/Clontech). All samples were sequenced in parallel at two different time points using Illumina NovaSeq 6000 flow cell and fastq files have been deposited on the Gene Expression Omnibus (GEO) database (GSE245307 and GSE268761 for males and females respectively). Reads were aligned to Ensembl GRCm39 annotations to quantify transcript abundance using Salmon 1.3.0 (Patro et al., 2017) and tximport (v1.18) R package (Soneson et al., 2016). For differential expression analysis, DESeq2 (v1.30) (Love et al., 2014) was performed in R environment (v4.0) using both gene and transcript level. Significance cut-off was set on a |log2fold change| > 0 and based on adjusted p-value < 0.1 for genes or adjusted p-value < 0.05 for transcripts. Finally, functional enrichment over representation analysis was performed using clusterProfiler (v3.18) Bioconductor package (Yu et al., 2012), in which statistically significance was defined based on adjusted p-value < 0.05 for each gene ontology enriched term.

### Reverse transcription

RNA concentration and integrity was measured using a NanoDrop spectrophotometer (Thermo Fisher Scientific). Reverse transcription was performed in similar amounts of RNA to produce cDNA in a 20 μl reaction using the High Capacity cDNA Reverse Transcription kit (Applied Biosystems) according to the manufacturer’s instructions.

### Quantitative real-time PCR analysis

Real-time PCR was carried out in a 10 μl reaction using 0.1 ng/μl of cDNA, using SYBR Green PCR Master Mix (Roche) according to the manufacturer’s protocol with a QuantStudio 12 K Flex Real-Time PCR System (Applied Biosystems).

Quantification was performed using the comparative CT Method (ΔΔCT Method). All the samples were tested in triplicate and the relative expression values were normalized to the expression value of β-actin. The fold change was calculated using the eq. 2(–ΔΔCt) formula, as previously reported (Livak and Schmittgen, 2001). The following primers specific for mouse were used:

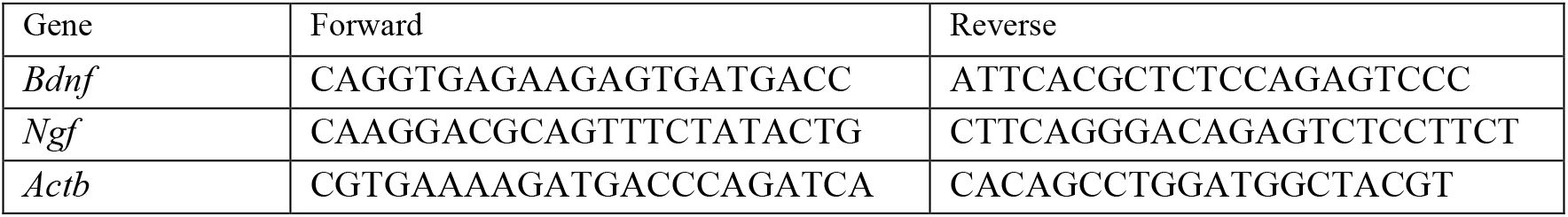

### Ex vivo electrophysiological recordings in the hippocampus

Hippocampal slices were prepared as described in detail elsewhere (Pérez-Rodríguez et al., 2019; Falcón-Moya et al., 2021). Briefly, mice were anesthetized with isoflurane (2% v/v) and brains were rapidly removed and placed into ice-cold solution (I) consisting of (in mM): 126 NaCl, 3 KCl, 1.25 KH2PO4, 2 MgSO4, 2 CaCl2, 26 NaHCO3, and 10 glucose (pH 7.2, 300 mOsmL-1), and positioned on the stage of a vibratome slicer and cut to obtain transverse hippocampal slices (350 mm thick), which were maintained continuously oxygenated for at least 1 h before use. All experiments were carried out at physiological temperature (30–34 °C). For experiments, slices were continuously perfused with the solution described above.

For LTP studies, whole-cell patch-clamp recordings were made from pyramidal cells located in the CA1 field of the hippocampus. CA1 pyramidal cells were patched under visual guidance by infrared differential interference contrast microscopy and verified to be pyramidal neurons by their characteristic voltage response to a current step protocol. Neurons were recorded using the whole-cell configuration of the patch-clamp technique in voltage-clamp mode with a patch-clamp amplifier (Multiclamp 700B), and data were acquired using pCLAMP 10.2 software (Molecular Devices). Patch electrodes were pulled from borosilicate glass tubing and had resistances of 4–7 MΩ when filled with (in mM): CsCl, 120; HEPES, 10; NaCl, 8; MgCl_2_, 1; CaCl_2_, 0.2; EGTA, 2 and QX-314, 20 (pH 7.2–7.3, 290 mOsm L−1). Cells were excluded from analysis if the series resistance changed by more than 15% during the recording.

In paired-pulse experiments, 2 consecutive stimuli separated by 40 ms were applied at the beginning of the baseline recording and again 30 min after applying the LTP protocol. Data were filtered at 3 kHz and acquired at 10 kHz. A stimulus-response curve (1–350 µA, mean of 5 excitatory postsynaptic current, EPSC, determination at each stimulation strength) was compiled for each experimental condition. Paired-pulse ratio (PRR) was expressed as the amplitude of the second response (EPSC) divided by the amplitude of the first (Falcón-Moya et al., 2021). Data were analysed using the Clampfit 10.2 software (Molecular Devices). The last 5 min of recording was used to estimate changes in synaptic efficacy compared to baseline.

### Tissue preparation for immunoblot analysis

For obtaining total hippocampal homogenates, hippocampal tissue was dounce-homogenized by 20 strokes with a loose pestle and 20 strokes with a tight pestle in 30 volumes of ice-cold lysis buffer (50 mM Tris-HCl pH 7.4, 150 mM NaCl, 10% glycerol, 1 mM EDTA, 1 μg/mL aprotinin, 1 μg/mL leupeptine, 1 μg/mL pepstatin, 1 mM phenylmethylsulfonyl fluoride, 1 mM sodium orthovanadate, 100 mM sodium fluoride, 5 mM sodium pyrophosphate, and 40 mM beta-glycerol phosphate) and 1% Triton X-100 using a Dounce homogenizer. After 10 min incubation at 4 °C, samples were centrifuged at 16,000 g for 30 min to remove insoluble fragments. Protein content in the supernatants was determined by DC-micro plate assay (Bio-Rad), following manufacturer’s instructions. Equal amounts of brain lysates were separated in 10% SDS-polyacrylamide gels before electrophoretic transfer onto nitrocellulose membrane overnight at 4 °C. Nitrocellulose membranes were blocked for 1 h at room temperature in Tris-buffered saline (TBS) (100 mM NaCl, 10 mM Tris, pH 7.4) with 0.1% Tween-20 (T-TBS) and 3% bovine serum albumin. Afterwards, nitrocellulose membranes were incubated for 2 h with the primary antibodies. Anti-BDNF (Icosagen, Cat# 327-100, RRID: AB_2927780) and anti-actin (Sigma-Aldrich Cat# MAB1501, RRID: AB_2223041) were used as primary antibodies. Then, nitrocellulose membranes were washed 3 times (5 min each) and subsequently incubated with the corresponding secondary antibody for 1 h at room temperature. Anti-mouse (Cell Signaling Technology Cat# 7076, RRID: AB_330924) was used as secondary antibody. After 3 washes (5 min each), immunoreactivity was visualized by enhanced chemiluminescence detection (Luminata Forte, Amersham). Optical densities of relevant immunoreactive bands were quantified after acquisition on a ChemiDoc XRS System (Bio-Rad) controlled by The Quantity One software v 4.6.9 (Bio-Rad). For quantitative purposes, the optical density values were normalized to β-actin values in the same sample and were expressed as a percentage of control treatment.

### Tissue preparation for immunofluorescence

Mice were deeply anesthetized by i.p. injection (0.2 mL/10 g of body weight) of a mixture of ketamine/xylazine (100 mg/kg and 20 mg/kg, respectively) prior to intracardiac perfusion of ice-cold 4% paraformaldehyde in 0.1 M phosphate buffer, pH7.4 (PB). Brains were removed and post-fixed overnight at 4 °C in the same fixative solution. The next day, brains were moved to 30% sucrose at 4 °C. Coronal brain sections of 30 μm were made on a freezing microtome (Leica SM 2000) and stored in a 5% sucrose solution at 4 °C until use.

### Ki67 immunofluorescence and cell quantification

Immunofluorescence and cell quantification were performed as previously described (Navarro-Romero et al., 2019). Brain sections were washed thrice with PB and blocked with 3% normal donkey serum and 0.3% Triton X-100 in PB (NDS-T-PB) at room temperature for 2 h. Slices were incubated overnight in the NDS-T-PB with the primary antibody to Ki67 (Abcam Cat# ab15580, RRID: AB_443209). The next day, sections were washed three times in PB and incubated at room temperature with the secondary antibody to rabbit immunoglobulins (Thermo Fisher Scientific Cat# A21206, RRID: AB_2535792) in NDS-T-PB for 2 h at room temperature. After incubation, sections were rinsed thrice with PB and then mounted onto glass slides coated with gelatine, in Fluoromount-G with DAPI (Thermo Fisher Scientific) as mounting medium.

Images of stained sections were obtained with a confocal microscope TCS SP5 LEICA (Leica Biosystems) using a dry objective (20×) with a sequential line scan at 1024 × 1024 pixel resolution. The images were obtained choosing a representative 10 μm Z-stack from the slice. The density of positive cells was quantified manually over the projection visualized after the application of an optimal automatic threshold (MaxEntropy) from Fiji software (ImageJ). The number of positive cells was calculated as the mean of total number of cells counted referred to the volume of the SGZ (μm3). Positive cells density was referred to that calculated for the control group.

### Statistical analysis

Results are reported as mean ± standard error of the mean (s.e.m). Data analyses were performed using GraphPad Prism software (GraphPad Prism 8). Statistical comparisons were evaluated using unpaired Student’s t-test for 2 groups comparisons or two-way ANOVA for multiple comparisons. Subsequent Bonferroni *post hoc* was used when required (significant interaction between factors). All results were expressed as mean ± s.e.m. Comparisons were considered statistically significant when *p* < 0.05.

## Results

### Repeated AM6545 treatment enhances novel object-recognition, place-recognition and fear memory

We first analysed the effect of a sub-chronic (7 days) treatment with AM6545 on memory performance. We tested novel object-recognition memory 48 h after training, as we previously demonstrated that at this time, control mice show reduced signs of object-recognition memory (Martínez-Torres et al., 2023). We observed that sub-chronic AM6545 treatment potentiated memory recall at this late time point in the NORT in both male (Student’s t-test: *p* = 0.002) (Figure 1A) and female (Student’s t-test: *p* = 0.001) mice (Figure 1B), similar to what we previously described with an acute administration (Martínez-Torres, 2023). These results with the sub-chronic administration indicate a lack of tolerance to the mnemonic effect of AM6545. We next performed the NORT under challenging conditions, by reducing the time of the training phase from 9 min to 3 min. Memory assessment 24 h later showed that AM6545 improved memory acquisition in both male (Student’s t-test: *p* = 0.0023) (Figure 1C) and female (Student’s t-test: *p* = 0.005) mice (Figure 1D), thus indicating that peripheral CB1R inhibition also enhances memory performance under challenging conditions suggesting a facilitation in memory consolidation. To assess the permanency of AM6545 effects in the NORT, we also performed the memory task one day after sub-chronic treatment withdrawal. Under these conditions, we still observed an enhancement in memory performance (Student’s t-test: *p* = 0.029) (Figure 1E), suggesting a permanent effect of repeated treatment with AM6545 over relevant cognitive brain areas. We have previously reported that pre-treatment with a single administration of the peripheral β-adrenergic blocker sotalol, prevents the mnemonic effect of acute AM6545 (Martínez-Torres et al., 2023). Here, we administered sotalol during 7 days 20 min before each administration of AM6545. We observed that sotalol pre-treatment totally prevented the pro-cognitive effects of AM6545 in the NORT (two-way ANOVA, interaction: F (1,52) = 7.993, *p* = 0.006; post hoc Bonferroni, Saline-VEH vs Saline-AM6545 *p* = 0.0014; Saline-AM6545 vs Sotalol-AM6545 *p* = 0.0018) (Figure 1F), indicating the participation of a peripheral adrenergic mechanism in the action of AM6545.

**Figure 1.**
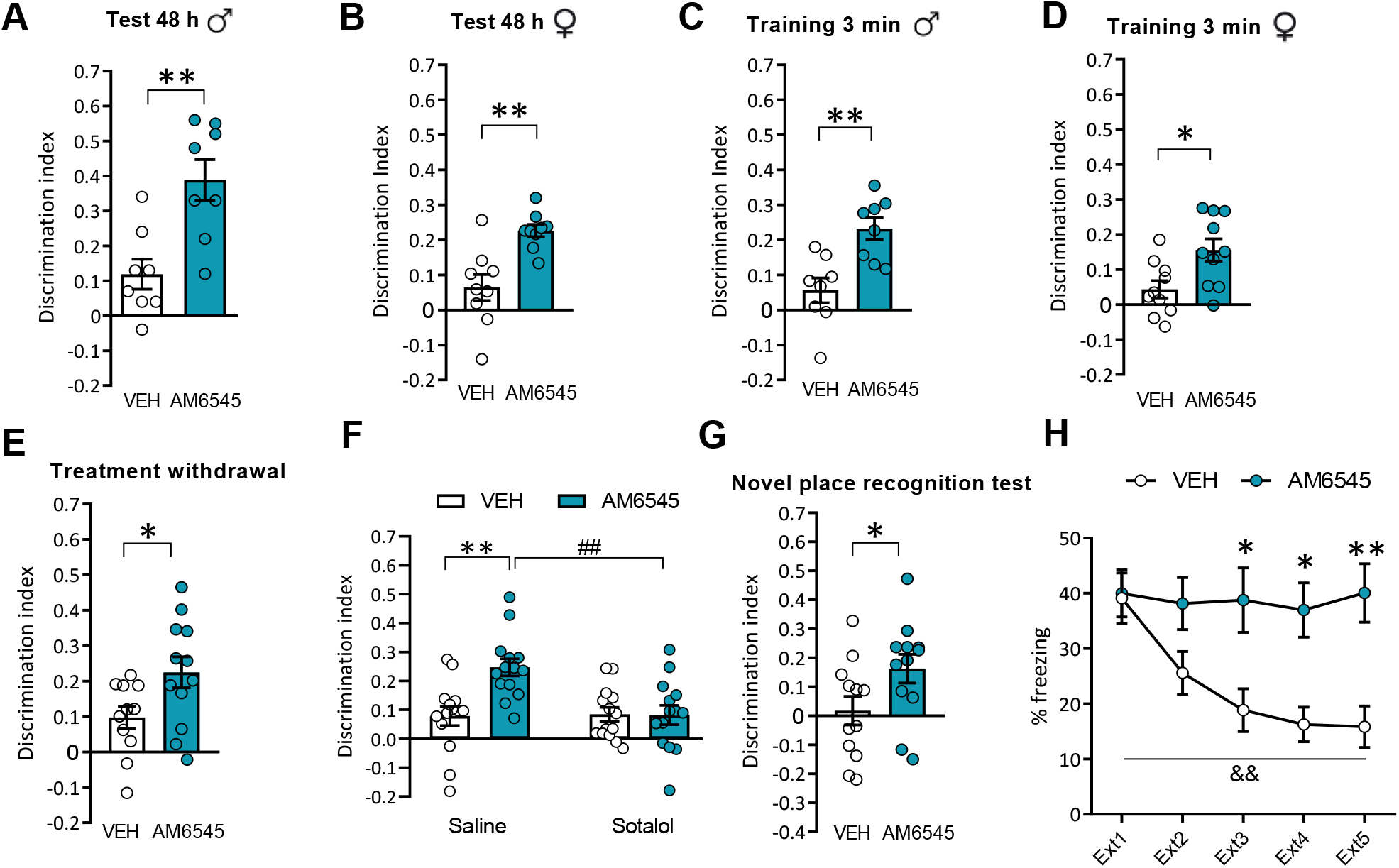
Sub-chronic AM6545 treatment improves both non-emotional and emotional memory. (**A,B)** Discrimination index values in NORT at 48 h after sub-chronic 7 days treatment with vehicle (VEH) or AM6545 (1 mg/kg) (*n* = 8-9) in both male (**A**) and female (**B**) mice. (**C,D)** Discrimination index values in NORT at 24 h after sub-chronic 7 d treatment with vehicle (VEH) or AM6545 (1 mg/kg) with a 3 min training period (*n* = 8-10) in both male **(C)** and female **(D)** mice**. (E)** Discrimination index values in NORT at 48 h after treatment withdrawal (*n* = 11-12). **(F)** Discrimination index values in NORT after sub-chronic 7 days treatment with vehicle (VEH) or AM6545 (1 mg/kg) after pre-treatment with saline or sotalol (10 mg/kg) (*n* = 14). **(G)** Discrimination index values in NPRT at 48 h after sub-chronic 7 days treatment with vehicle (VEH) or AM6545 (1 mg/kg) (*n* = 12). **(H)** Percentage of freezing in the context fear conditioning across extinction sessions (Ext1-Ext5) after post-extinction 1-5 treatment with vehicle (VEH) or AM6545 (1 mg/kg) (*n* = 16). Data are expressed as mean ± s.e.m. * *p* < 0.05, ** *p* < 0.01 (treatment effect), ## *p* < 0.01 (pre-treatment effect), && *p* < 0.01 (time effect) by Student’s t-test or two-way repeated measures ANOVA followed by Bonferroni *post hoc*.

Additionally, we assessed if the repeated exposure to AM6545 was able to improve other memory domains such as spatial memory using the novel place-recognition test (NPRT). When we tested memory at 48 h after the training phase, the sub-chronic AM6545 treatment showed an enhancement in spatial memory persistence (Student’s t-test: *p* = 0.049) (Figure 1G), pointing to a role in modulating also spatial memory. In order to discard anxiety-related effects, as has been described previously for the systemic inhibition of CB1R in rodents (Patel and Hillard, 2006), we evaluated the effect of the peripheral CB1R antagonist on the elevated plus-maze test after 7 days of treatment. We observed that the percentage of time and entries to the open arms did not significantly change (Supplementary Fig. 1A-B) compared to vehicle-treated mice, indicating that AM6545 at a dose of 1 mg/kg is not producing an unspecific effect in the anxiety-like response. We also assessed locomotor activity in actimeter boxes after sub-chronic administration of the peripheral CB1R antagonist AM6545 and found no significant effect of AM6545 in the locomotor activity as reflected in total activity and rearings measurements (Supplementary Fig. 1C-E).

Next, we studied whether peripheral CB1R inhibition would modify memory persistence in an emotional task using the context fear conditioning paradigm, by administering AM6545 (1 mg/kg, i.p.) after each extinction session. We observed that AM6545-treated mice prevented fear extinction across extinction sessions (Ext2-Ext5) (two-way repeated measures ANOVA, treatment effect: F (4,75) = 3.178, *p* = 0.018; post hoc Bonferroni, Ext3-VEH vs Ext3-AM6545, *p* = 0.024, Ext4-VEH vs Ext4-AM6545, *p* = 0.017, Ext5-VEH vs Ext5-AM6545, *p* = 0.003) (Figure 1H).

### Long-term treatment with AM6545 improves executive function in the pairwise discrimination task

To test whether the peripheral CB1R inhibition modifies other cognitive domains, we explored executive function by performing the pairwise discrimination task using the touchscreen chambers with previously validated images (Figure 2A) (Boyers et al., personal communication). We administered AM6545 every day after the pairwise discrimination session until mice completed the task. We observed that AM6545-treated mice showed a more rapid learning compared to vehicle as reflected in a smaller number of correction trials (Student’s t test: *p* = 0.001), and a reduced number of sessions to complete the task (Student’s t test: *p* = 0.026) (Figure 2B-C). There was a non-significant (Student’s t-test: *p* = 0.10) trend for AM6545-treated mice to perform less trials to reach criterion (Figure 2D). We also analysed other control parameters such as the reward collection latency, the correct and incorrect touch latency, the left and right intertrial interval (ITI) touches and the time to initiate the trials. No significant differences in any of these parameters were observed (Supplementary Fig. 2A-F). There were also no significant changes in the body weight variation (Δbody weight) between groups during the protocol (Supplementary Fig. 2G), altogether indicating no major changes in general activity produced by AM6545, further supporting the relevance of its effect on cognitive performance.

**Figure 2.**
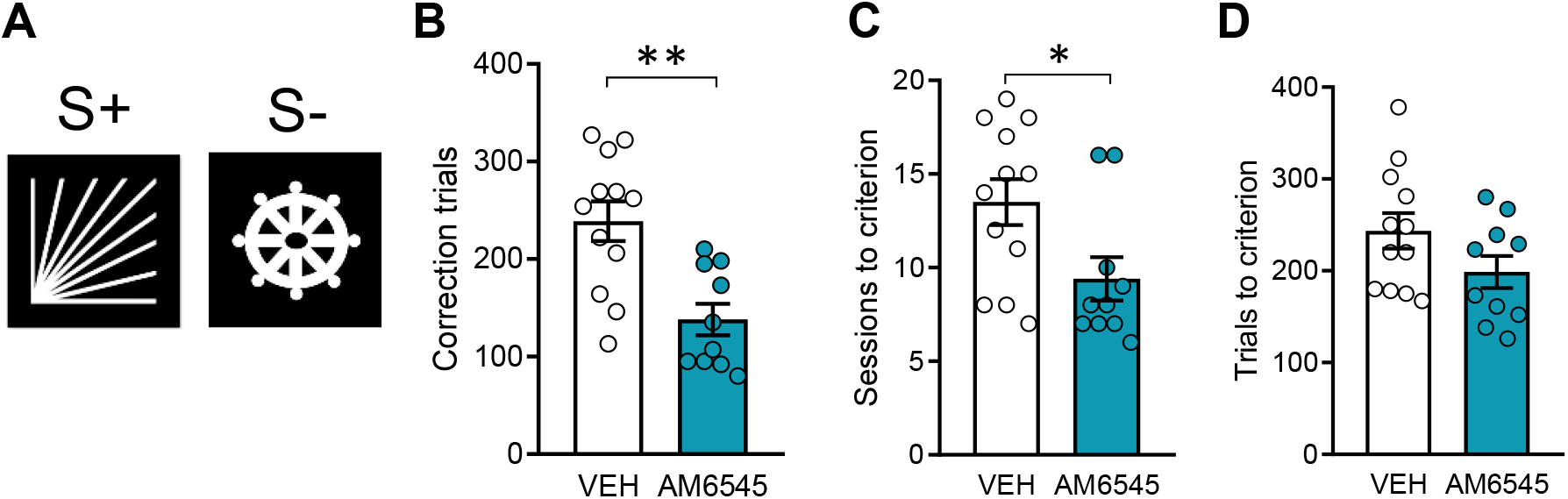
Long-term peripheral CB1R inhibition improves executive function in the pairwise discrimination task. (**A)** Image pair presented in the touchscreen during the pairwise discrimination test. (**B-D)** Total number of correction trials **(B)**, sessions **(C)** and trials **(D)** required to reach criterion in animals treated with vehicle (VEH) or AM6545 (1 mg/kg) (*n* = 10-12). Data are expressed as mean ± s.e.m. * *p* < 0.05, ** *p* < 0.01 by Student’s t-test.

### Peripheral CB1R inhibition differentially modulates synaptic transcriptome in male and female mice

We studied the potential synaptic modifications associated with the pro-cognitive effect produced by AM6545 in both male and female mice. For this, we used high throughput RNA sequencing to analyse transcriptomic changes between five male mice and three female mice sub-chronically treated with vehicle and five male mice and four female mice sub-chronically treated with AM6545 (1 mg/kg, i.p.). Hippocampal tissues were obtained 1 day after treatment withdrawal. As we wanted to evaluate synaptic modifications promoted by peripheral CB1R modulation, we focused on hippocampal synaptoneurosomal enriched fractions using a subcellular enrichment previously characterized in our group (Salgado-Mendialdúa et al., 2018). When we analysed gene expression, few genes appeared differentially expressed in male mice (Supplementary Fig. 3 and Supplementary Table 1). Surprisingly, in female mice we found the upregulation of 18 genes and the downregulation of 113 genes (Supplementary Fig. 3 and Supplementary Table 1).

Assuming that synaptic contacts are fractions with local translation and with high abundance in transcript content, we also analysed differential expression at transcript level (Hafner et al., 2019). We found that 278 transcripts were differentially modulated in AM6545 treated male mice compared with vehicle (Figure 3A). In fact, with an adjusted p-value < 0.05, 133 transcripts were upregulated while 145 transcripts were downregulated (VEH-male *vs.* AM6545-male). Regarding gene ontology analysis, we observed that upregulated transcripts present an enrichment in terms related with synapse structure and activity and pre- and post-density localization (Figure 3B). However, downregulated transcripts were less associated between them as we obtained more specific molecular functions such as methyltransferase activity or catalytic processes and also some related with synapse development (Figure 3B). In the case of female mice, 580 transcripts were differentially expressed due to AM6545 administration (Figure 3C). Specifically, we found the upregulation of 345 transcripts and the downregulation of 235 (VEH-female *vs.* AM6545-female). In terms of gene ontology enrichment, we found that upregulated transcripts were associated mainly with structural synapse and postsynaptic density although we also found transcripts related with histone and protein modification (Figure 3D), processes that can be related to the required protein synthesis that change synaptic plasticity. In contrast, most significant terms associated with downregulated transcripts in female mice can be summarized as mitochondrial and ribosomal processes (Figure 3D) and similarly as with males, we also found some enriched terms related to synapse development.

**Figure 3.**
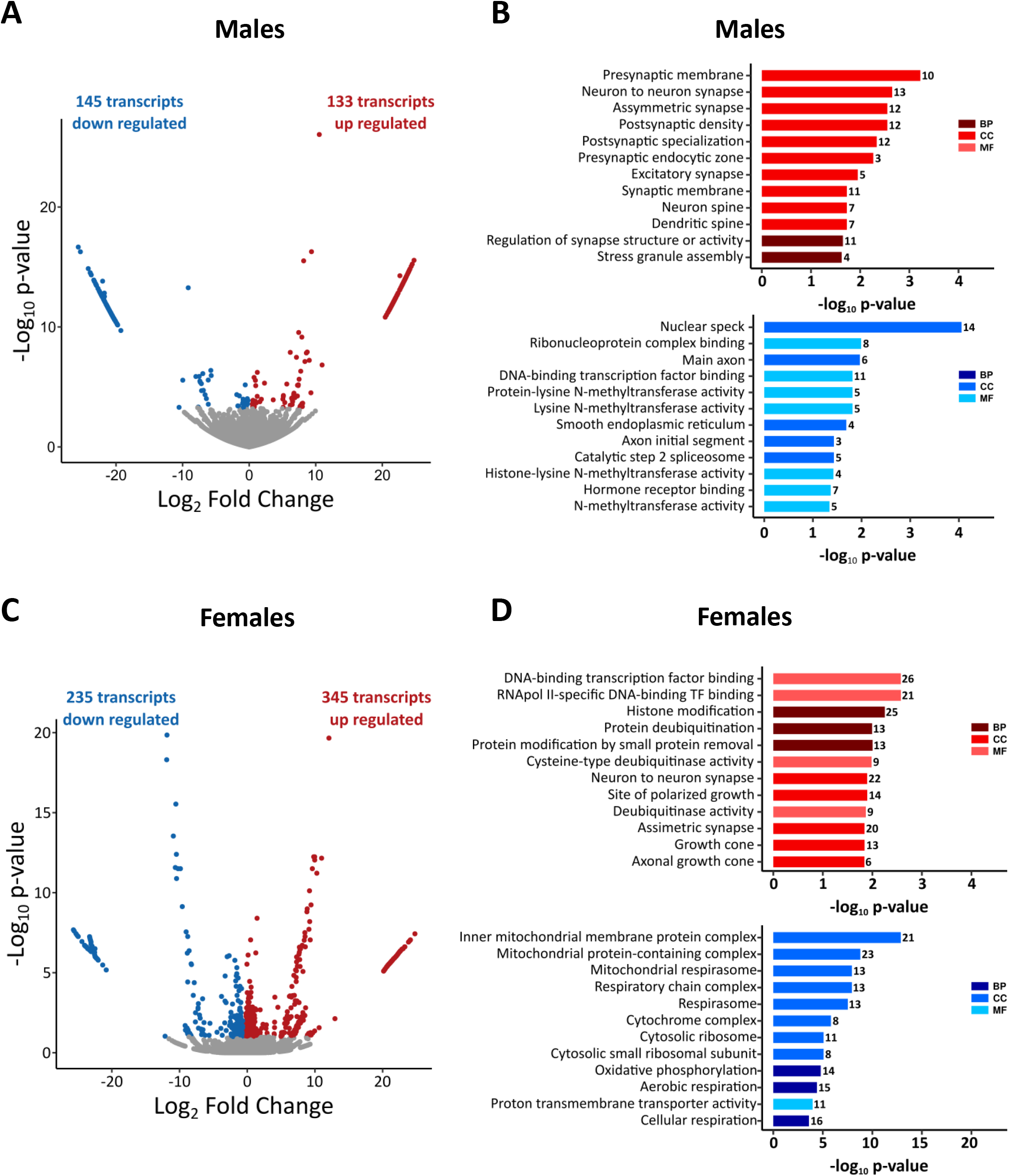
Sub-chronic peripheral CB1R inhibition differentially modulates male and female hippocampal transcript expression. **(A)** Volcano plot of differentially expressed transcripts between vehicle and AM6545-treated male mice. **(B)** Gene ontology enrichment analysis for both upregulated and downregulated genes associated to each transcript in AM6545-treated male mice compared to vehicle. **(C)** Volcano plot of differentially expressed transcripts between vehicle and AM6545-treated female mice. **(D)** Gene ontology enrichment analysis for both upregulated and downregulated genes associated to each transcript in AM6545-treated female mice compared to vehicle. In volcano plots, red indicates significant upregulation (p < 0.05 and log2FC > 0) and blue significant downregulation (p < 0.05 and log2FC < 0). Most significant GO terms are represented, red and blue indicate upregulation or downregulation respectively and the intensity indicates GO category of each term. (VEH-male, n = 5; AM6545-male, n = 5; VEH-female, n = 3; AM6545-female, n = 4).

### Sub-chronic AM6545 treatment induces synaptic plasticity changes in the hippocampus

We first assessed cellular plasticity by examining the proliferation of neuronal precursors in the hippocampus. Adult neurogenesis was studied through the quantification of the number of cells expressing the endogenous marker of cell proliferation Ki67 in the subgranular zone (SGZ) of the dentate gyrus. Sub-chronic treatment with AM6545 did not significantly modify the number of Ki67+ cells, suggesting that AM6545 does not modify hippocampal neurogenesis (Supplementary Fig. 4).

Next, to investigate the functional consequences of peripheral CB1R inhibition in hippocampal plasticity, we studied long-term potentiation (LTP) at Schaffer collateral-CA1 synapses in slices from mice treated for 7 days with AM6545 (1 mg/kg) or vehicle. Acute hippocampal slices were obtained 24 h after the last drug administration, the time when novel object-recognition memory assessment, transcriptomic analyses and Ki67 analyses were performed. LTP was induced by stimulating Schaffer collaterals at 100-Hz during 1s. Slices from vehicle-treated mice showed robust LTP (168 ± 10 %), whereas LTP was completely occluded in slices from mice treated with AM6545 (96 ± 8 %) (Student’s t-test: *p* = 0.002) (Figure 4A-B). To determine the site of expression of LTP, we analysed pair-pulse ratios (PPR) during baseline and 30 min after LTP induction. This analysis showed no differences in control slices (1.41 ± 0.11 after LTP vs 1.67 ± 0.16 in baseline) (Figure 4C) confirming the established postsynaptic expression of this form of LTP. Repeated treatment with AM6545 did not induce changes in PPR (1.51 ± 0.18 after LTP vs 1.89 ± 0.17 in baseline). We also compiled a stimulus-response curve (50-350 µA) to examine whether the basal synaptic transmission would be modified in mice treated with AM6545 for 7 days. We found that slices from AM6545-treated mice presented increased amplitude of EPSCs (Figure 4D), which could explain the LTP occlusion described above.

**Figure 4.**
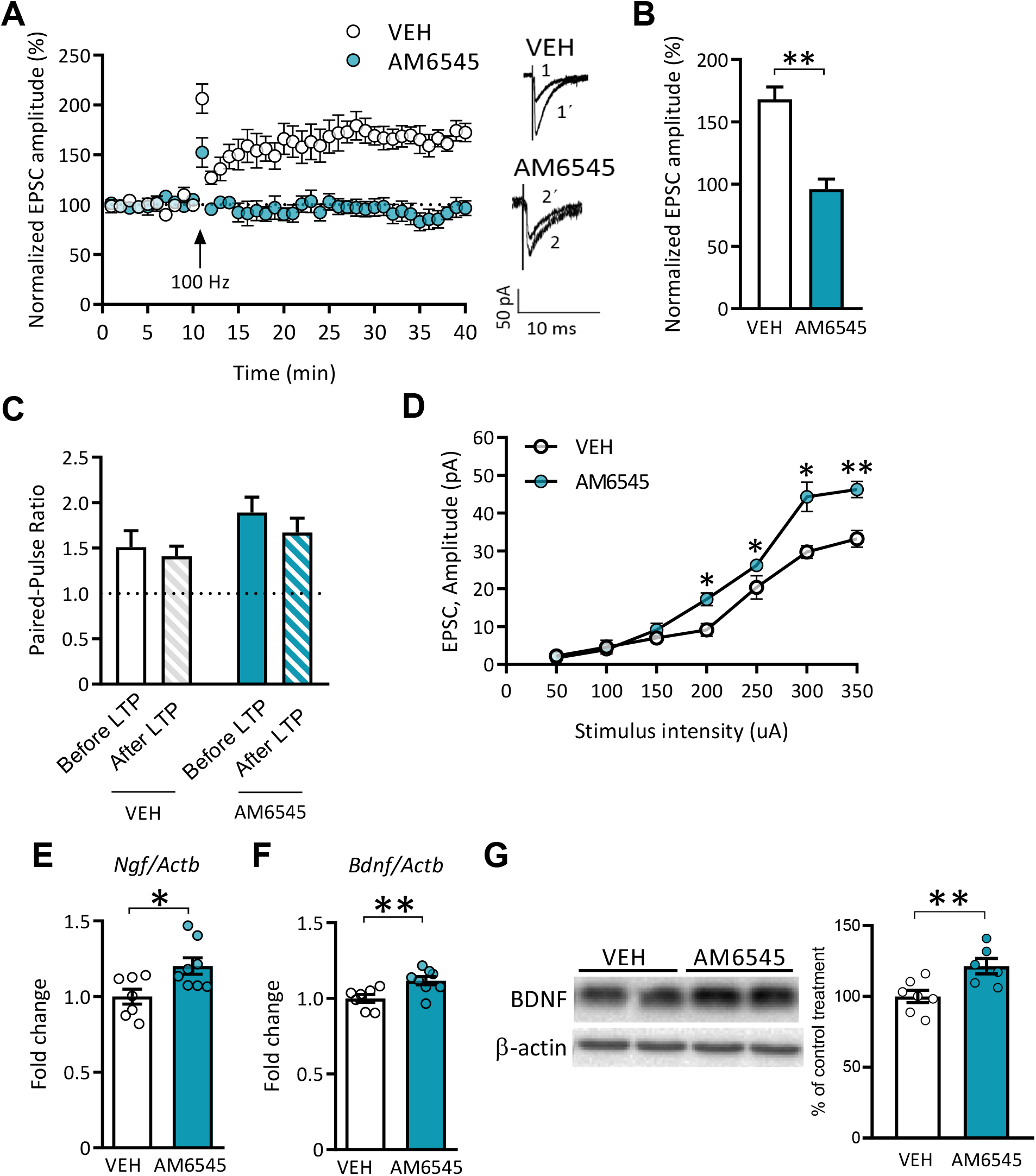
Sub-chronic peripheral CB1R inhibition modifies synaptic plasticity in the hippocampus. **(A)** Average time courses of the change in the amplitude of the EPSC in hippocampal slices from mice treated for 7 days with vehicle (VEH) or AM6545 (1 mg/kg). Traces represent samples of EPSC recorded for each experimental group before (1,2) and 30 min after (1’,2’) LTP induction (*n* = 6). **(B)** Average LTP of the last 5 min of recordings (*n* = 6). (**C)** Paired-pulse ratio before and after (fill pattern bars) LTP induction in hippocampal slices from male mice treated for 7 days with vehicle (VEH) or AM6545 (1 mg/kg). (**D)** Stimulation input/output curves in hippocampal slices from mice treated for 7 days with vehicle (VEH) or AM6545 (1 mg/kg). **(E,F)** Hippocampal mRNA levels of the neurotrophic factors *Ngf* **(E)** and *Bdnf* **(F)** of mice treated for 7 days with vehicle (VEH) or AM6545 (1 mg/kg) (n = 7 - 8). **(G)** Representative immunoblot and quantification of BDNF total expression in the hippocampus of mice treated for 7 days with vehicle (VEH) or AM6545 (1 mg/kg) (n = 7). Data are expressed as mean ± s.e.m. * *p* < 0.05, ** *p* < 0.01 by Student’s t-test or two-way repeated measures ANOVA followed by Bonferroni *post hoc*.

Finally, we evaluated the expression of neurotrophic factors in the hippocampus of male mice to assess the mechanism involved in AM6545 effect on synaptic plasticity. Quantitative RT-PCR analysis showed a significant enhancement in the mRNA levels of *Ngf* (Student’s t-test: *p* = 0.018) (Figure 4E) and *Bdnf* (Student’s t-test: *p* = 0.009) (Figure 4F) after sub-chronic 7 days AM6545 treatment. Interestingly, immunoblot analysis supported the increase of BDNF levels in hippocampal homogenates of mice treated for 7 days with AM6545 (Student’s t-test: *p* = 0.009) (Figure 4G).

## Discussion

In this study we reveal a relevant effect of peripheral CB1R inhibition in enhancing memory and executive function in mice. Such cognitive improvements were accompanied by differential hippocampal transcript expression and modifications in hippocampal synaptic plasticity and hippocampal neurotrophin expression.

Previous studies showed that systemic CB1R antagonists could improve memory (Terranova et al., 1996; Wolff and Leander, 2003; Takahashi et al., 2005). In addition, genetic downregulation of CB1R also enhances memory performance (Reibaud et al., 1999; Maccarrone et al., 2002; Martínez-Torres et al., 2023). In this study we used the peripherally-restricted CB1R antagonist AM6545. Our previous study proved that an acute administration of AM6545 produced improvements in novel object-recognition memory. Despite the high affinity of AM6545 for mouse and rat CB1R, previous data has suggested that AM6545 may affect the hypothalamic-pituitary-adrenal axis via a non-CB1R mechanism (Roberts and Hillard, 2020). However, the doses used in this study were 5 and 10 times higher than the dose used in the current study. Here, we used the dose of 1 mg/kg as we previously demonstrated that this dose was optimal for the pro-cognitive effects of AM6545, and that lower and higher doses did not show the mnemonic outcomes. Moreover, we also observed that another peripherally-restricted CB1R antagonist, TM38837 also enhanced recognition memory persistence, thus supporting a relevant CB1R-dependent mechanism independent of the specific drug used (Martínez-Torres et al., 2023). All this evidence showed that peripheral CB1R could be a potential target in memory modulation. In the present study, we used a sub-chronic 7 days administration of the peripherally-restricted CB1R antagonist AM6545 as a strategy to explore additional cognitive domains sensitive to CB1R peripheral inhibition. Here, we demonstrated that no tolerance was developed after repeated AM6545 treatment since treatment efficacy was observed after 7 administrations. Indeed, object-recognition memory persistence was significantly enhanced in both male and female mice when the test was performed 48 h after training at the end of treatment. In this case, we used the period of 48 h as we previously revealed that novel object-recognition memory is a transient memory trace and at the 48 h time point, control mice naturally show reduced novel object discrimination and hence low discrimination indexes (Martínez-Torres et al., 2023). Moreover, we also observed that the repeated AM6545 treatment improved memory in both sexes when NORT was performed under challenging conditions by reducing the training period to 3 min, compared to the 9 min in standard conditions. Under these conditions AM6545-treated mice exhibited higher discrimination indexes than control mice, indicating that the treatment enhances memory facilitation. Similar to our approach, the reduction of the training period in the NORT has been previously used to assess memory enhancement through exercise in mice (Butler et al., 2019). Additionally, we assessed recognition memory in the NORT one day after treatment withdrawal, thus the drug was not onboard during memory consolidation. Such conditions also enhanced memory persistence, suggesting that memory improvement could be mediated by persistent brain plasticity changes elicited by repeated treatment.

Our previous research showed that single administration AM6545-memory improvement was dependent on a peripheral adrenergic mechanism involving the adrenal glands given that AM6545 mnemonic effect was prevented by the peripherally restricted β-adrenergic antagonist sotalol and was absent in adrenalectomized mice (Martínez-Torres et al., 2023). In this regard, the fact that here sotalol pre-treatment significantly reduced the mnemonic effect of sub-chronic AM6545 points to a relevant role of the peripheral adrenergic tone. This is congruent with the fact that the release of adrenaline and noradrenaline from the adrenal glands has a significant effect on memory persistence (McIntyre et al., 2012; Yang et al., 2013). In addition, peripheral adrenergic receptor inhibition prevented the improvement in memory persistence elicited by novelty in a fear conditioning paradigm (King and Williams, 2009), further supporting a peripheral adrenergic mechanism involved in memory consolidation where we propose peripheral CB1R could participate.

We also assessed whether other memory domains could be sensitive to this pharmacological intervention with AM6545. We found a significant enhancement in the NPRT, a type of spatial memory. This effect is reminiscent of previous results where systemic CB1R inhibition with the CB1R antagonist AM281 improved spatial learning in the Morris water maze test in mice with traumatic brain injury (Xu et al., 2019). To the best of our knowledge, the results reported herein are the first to point to a potential role of the peripheral CB1R in spatial memory performance in young adult mice.

Regarding emotional memory, we previously reported that a single administration of AM6545 after the first extinction session in the context fear conditioning, promoted freezing behaviour when memory was assessed the next day. However, freezing was reduced when assessed in the subsequent extinction sessions without AM6545 re-exposure (Martínez-Torres et al., 2023). In the present study, we demonstrate that administering AM6545 after every extinction session maintains high freezing levels across days, pointing to a persistent effect of repeated exposure to AM6545 in emotional memory reconsolidation. This finding is reminiscent of the results obtained with systemic approaches such as the use of the CB1R antagonist rimonabant or the use of the constitutive CB1R KO mice (Marsicano et al., 2002), and point again to the periphery for effects assumed to depend on central inhibition of CB1R.

Remarkably, when we assessed executive function, we found that mice that received repeated AM6545 administration had a better performance in the pairwise discrimination task. In this regard, CB1R function seems relevant for this type of cognitive task, since the use of a systemic CB1R agonist impaired performance in another executive function-related task whereas the administration of a systemic CB1R antagonist prevented this impairment (Arguello and Jentsch, 2004). Here, we report that the administration of the peripherally-restricted CB1R antagonist after each training session improves executive function performance, as reflected in the number of correction trials and sessions to criterion.

The transcriptomic analysis in hippocampal synaptoneurosomal fractions showed different effects in AM6545-treated male and female mice. Male mice showed very limited number of changes at the gene expression level, while female mice showed the upregulation of genes potentially related with the pro-cognitive effects of AM6545. We identified *Hap1*, a gene associated with BDNF signalling (Lim et al., 2018) or Pik3ca, associated to the PI3K-AKT-mTOR pathway which controls translation (Borrie et al., 2017). Among the downregulated genes in females, we found members of *Rpl* and *Rps* ribosomal families which may also indicate an optimization of ribosomal composition to improve activity-induced translation at synapses (Beletskiy et al., 2024). When all possible transcripts detected in the high-throughput study were considered independently of the gene transcribed, more specific modifications between treatment conditions were revealed both in male and female mice, allowing the implementation of an enrichment analysis. Interestingly, we found that gene ontology terms associated with synapse were in common among upregulated and downregulated transcripts in both sexes, pointing to prominent effects of peripheral CB1R inhibition on the synaptic transcriptome at the level of splice variants. Further analyses will be needed to establish causal relationship in the possible sexual dimorphic mechanisms of AM6545 in memory modulation.

We also evaluated neuronal progenitor proliferation in the subgranular zone of the dentate gyrus. Previous evidence linked adult neurogenesis in this hippocampal subregion to the establishment of hippocampal-dependent memory (Saxe et al., 2006; Dupret et al., 2008; Jessberger et al., 2009). However, AM6545 treatment did not modify the number of cells expressing the cell proliferation marker Ki67 in this brain region, indicating that AM6545 did not affect this process. Instead, our electrophysiological studies confirmed that sub-chronic AM6545 treatment increased the amplitude of EPSCs in the hippocampal CA1 region. As the experiments were performed at -70 mV, EPSCs were mediated by the activation of AMPA receptors. No changes in PPR were observed after AM6545 treatment, mainly discarding a presynaptic component in the facilitation of neurotransmitter release, and suggesting a postsynaptic increase in AMPA receptor-mediated currents. In the hippocampus, the activation of β-adrenergic receptors, which could result from AM6545 administration, according to its described acute effects (Martinez-Torres et al., 2023), can trigger the phosphorylation of AMPAR facilitating their traffic to extrasynaptic sites (Vanhoose and Winder, 2003; Rouach et al., 2005; Joiner et al., 2010), reinforcing LTP (Oh et al., 2006) by reducing the threshold for induction (Papaleonidopoulos and Papatheodoropoulos, 2018), and potentially leading to saturation and occlusion, an explanation parsimonious with our experimental observations.

The neuronal plasticity changes elicited by AM6545 repeated administration could be also mediated by the increase of mRNA level of *Bdnf* and *Ngf* in the hippocampus of AM6545-treated mice and a concomitant increase of BDNF protein levels. These neurotrophins have an important role in long-term synaptic plasticity and memory (Gibon and Barker, 2017) and may contribute to the overall pro-cognitive effects of AM6545.

Altogether, our study suggests that sub-chronic peripheral CB1R inhibition contributes to the enhanced performance of low arousal and emotional memory persistence as well as executive function. These behavioural outcomes may be mediated by different hippocampal synaptic plasticity changes, mimicking the effects of systemic approaches reducing CB1R function and pointing to peripheral CB1R as potential targets to improve cognitive performance.

## Acknowledgements

We are grateful to Marta Linares, Raquel Martín and Francisco Porrón for expert technical assistance and Lorena Galera-López, Laura Ciaran-Alfano and Hira Nizam for helpful discussion.

## Author contributions

A.B.-M.: Behavioural and biochemical experiments, statistical analyses and graphs and writing of the manuscript; L.R.-R.: Transcriptomic study; S.M.-T.: Behavioural experiments; I.M.-G. and A.R.-M.: Electrophysiological experiments; R.M.: Supervision and funding of the study; A.O.: Conceptualization, supervision, funding of the study, and writing of the manuscript. All authors revised the final version of the manuscript.

## Supporting Information

**Supplementary Figure 1.**
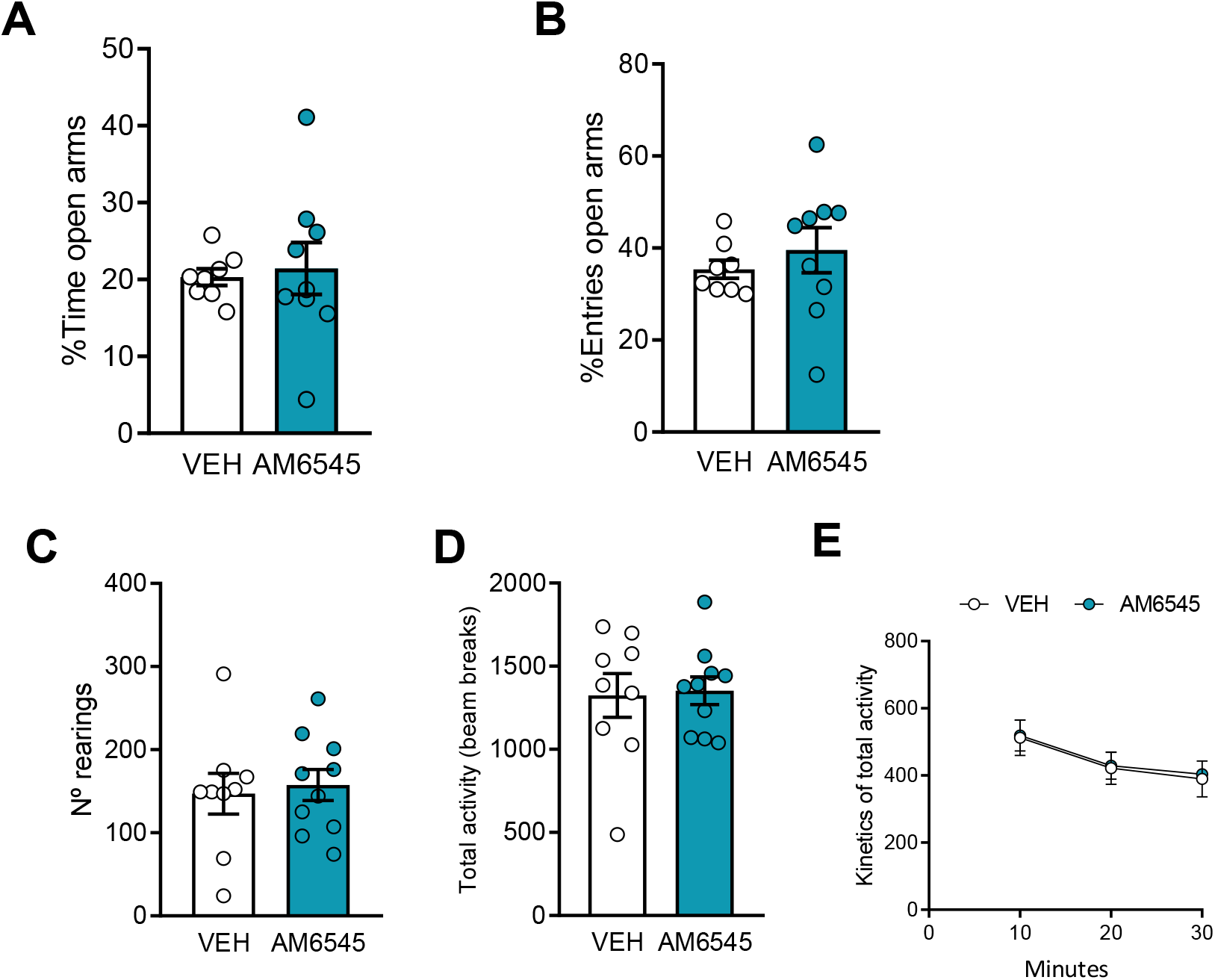
Sub-chronic AM6545 treatment does not modify anxiety-like behavior and locomotor activity. **(A,B)** Percentage of time **(A)** and entries **(B)** to the open arms in the elevated-plus maze test after sub-chronic 7 days treatment with vehicle (VEH) or AM6545 (1 mg/kg) (*n* = 8 – 9). **(C-E)** Total number of rearings **(C)**, total activity **(D)** and kinetics of total activity **(E)** during 30 min after sub-chronic 7 days treatment with vehicle (VEH) or AM6545 (1 mg/kg) (*n* = 9 – 10). Data are expressed as mean ± s.e.m. Statistical significance was calculated by Student’s t test or two-way repeated measures ANOVA.

**Supplementary Figure 2.**
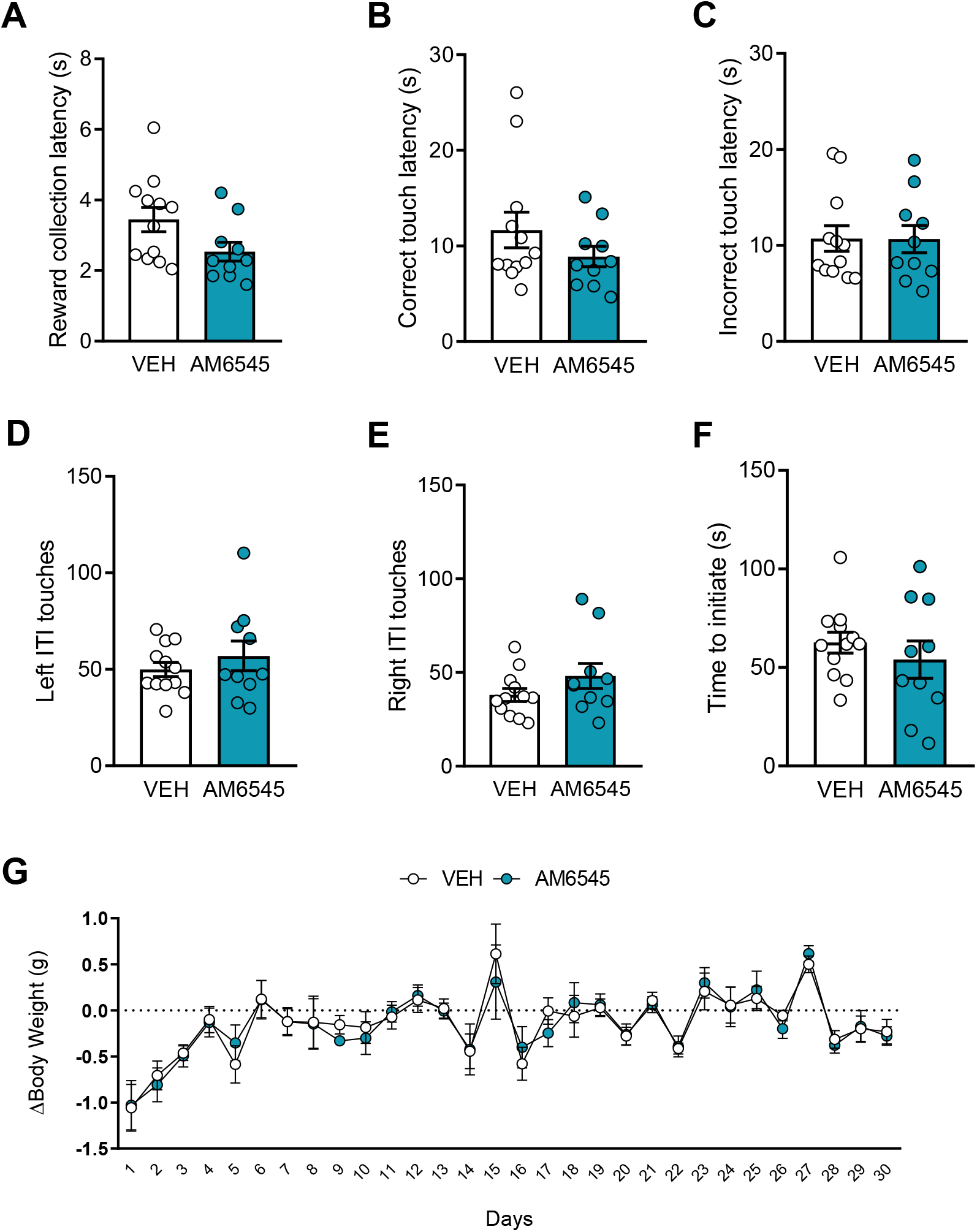
Long-term peripheral CB1R inhibition does not modify control parameters or body weight in the pairwise discrimination task. **(A-F)** Reward collection latency (**A**), correct **(B)** and incorrect (**C**) touch latency, left (**D**) and right (**E**) ITI touches and time to initiate the trials (**F**) in animals treated with vehicle (VEH) or AM6545 (1 mg/kg) (*n* = 12-14). (**G**) Weight variation between days in animals treated with vehicle (VEH) or AM6545 (1 mg/kg) (*n* = 12-14). Data are expressed as mean ± s.e.m. Statistical significance was calculated by Student’s t-test or two-way ANOVA.

**Supplementary Figure 3.**
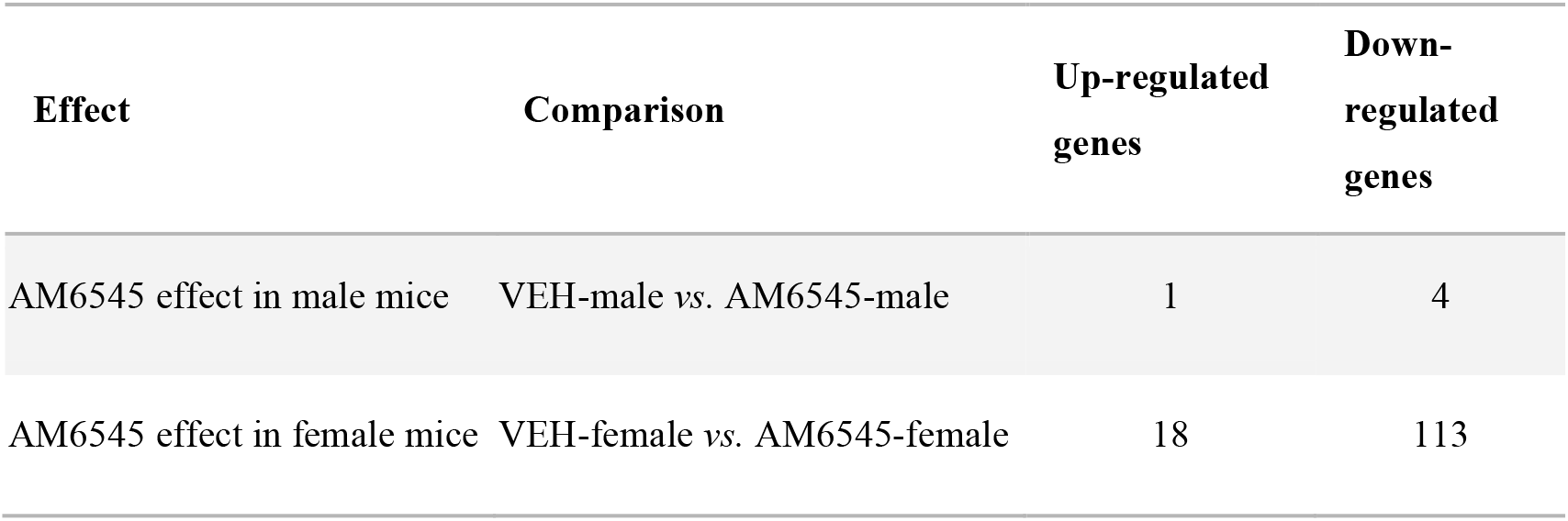
Sub-chronic AM6545 administration modifies more genes in female than male hippocampal synaptoneurosomes. Summary of differentially expressed genes in hippocampal synaptoneurosomes from mice treated with AM6545 compared to mice treated with vehicle in both male and female mice. Threshold for significance was settled at adjusted p-value < 0.1. (VEH-male, *n* = 5; AM6545-male, *n* = 5; VEH-female, *n* = 3; AM6545-female, *n* = 4). Statistical significance was calculated by Benjamini-Hochberg adjustment following Wald test analysis.

**Supplementary Figure 4.**
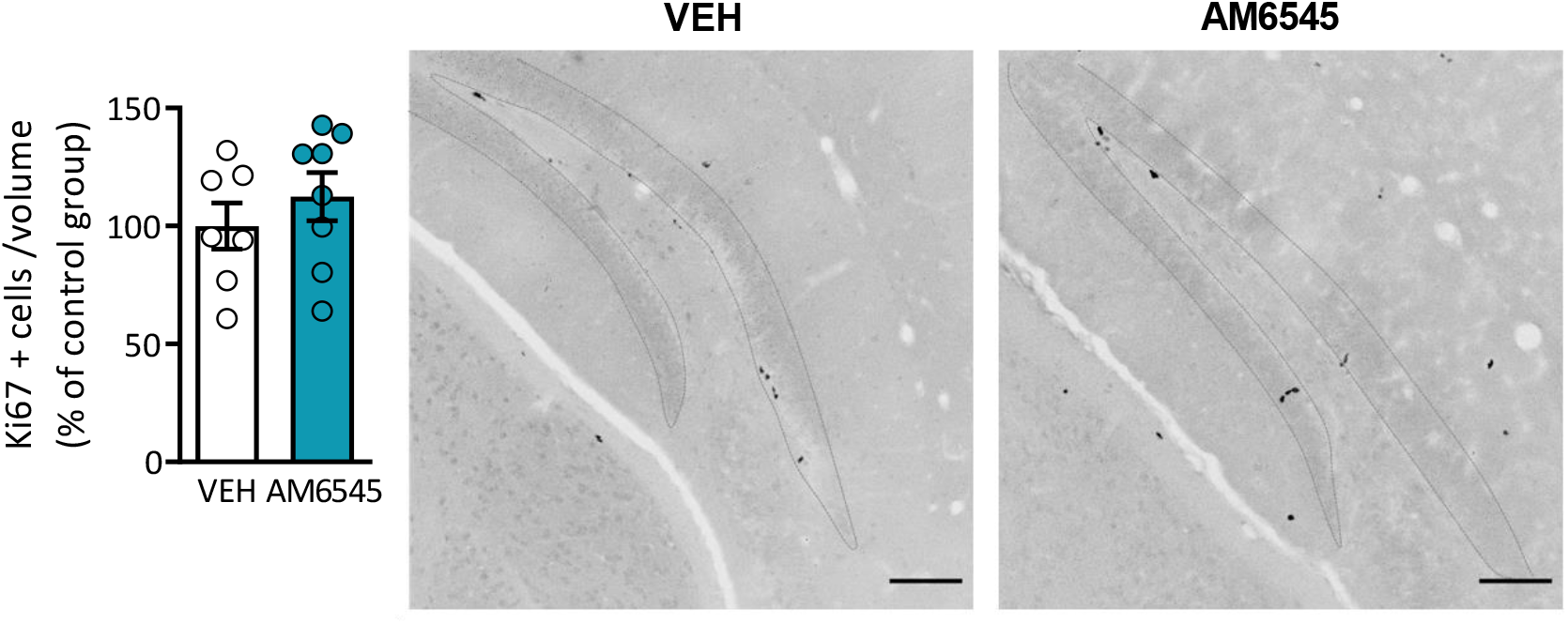
Sub-chronic peripheral CB1R inhibition does not modify hippocampal neurogenesis. Average density of Ki67+ cells and representative greyscale confocal images in the subgranular zone of the dentate gyrus of mice treated for 7 days with vehicle (VEH) AM6545 (1 mg/kg) (*n* = 7 - 8) (scale bar = 100 μm). Data are expressed as mean ± s.e.m. Statistical significance was calculated by Student’s t-test.

